# Revisiting the role of *Toxoplasma gondii* ERK7 in the maintenance and stability of the apical complex

**DOI:** 10.1101/2021.03.22.436543

**Authors:** Nicolas Dos Santos Pacheco, Nicolò Tosetti, Aarti Krishnan, Romuald Haase, Dominique Soldati-Favre

## Abstract

*Toxoplasma gondii* ERK7 is known to contribute to the integrity of the apical complex and to be involved only in the final step of the conoid biogenesis. In the absence of ERK7, mature parasites lose their conoid complex and are unable to glide, invade or egress from host cells. In contrast to a previous report, we show here that depletion of ERK7 phenocopies the depletion of the apical cap proteins AC9 or AC10. The absence of ERK7 leads to the loss of the apical polar ring, the disorganization of the basket of subpellicular microtubules and an impairment in micronemes secretion. Ultra-expansion microscopy (U-ExM) coupled to NHS-Ester staining on intracellular parasites offers an unprecedented level of resolution and highlights the disorganization of the rhoptries as well as the dilated plasma membrane at the apical pole in the absence of ERK7. Comparative proteomics analysis of wild-type and ERK7 or AC9 depleted parasites led to the disappearance of known, predicted, as well as putative novel components of the apical complex. In contrast, the absence of ERK7 led to an accumulation of microneme proteins, resulting from the defect in exocytosis of the organelles.

**Importance:** The conoid is an enigmatic, dynamic organelle positioned at the apical tip of the coccidian subgroup of the Apicomplexa, close to the apical polar ring (APR) from which the subpellicular microtubules (SPTMs) emerge and at the site of microneme and rhoptry secretory organelles exocytosis. In *Toxoplasma gondii*, the conoid protrudes concomitantly to microneme secretion, during egress, motility and invasion. The conditional depletion of the apical cap structural protein AC9 or AC10 leads to a disorganization of SPTMs as well as the loss of APR and conoid that result in microneme secretion defect and block in motility, invasion and egress. We show here that depletion of the kinase ERK7 phenocopies completely AC9 and AC10 mutants. Moreover, the combination of ultrastructure expansion microscopy with an NHS ester staining revealed that ERK7 depleted parasites exhibit a dilated apical plasma membrane and a mis-positioning of the rhoptry organelles.

## Introduction

The Apicomplexa phylum is defined by the so-called apical complex, a structure that harbors unique secretory organelles, termed rhoptries and micronemes, as well as membranous and cytoskeletal elements that critically contributes to parasite dissemination. The overall cytoskeleton is made of three interacting layers, conferring structure and stability to the parasite: the inner membrane complex (IMC), formed by a patchwork of flattened vesicles, the subpellicular network (SPN) composed of intermediate filaments-like proteins termed alveolins and the subpellicular microtubules (SPMTs). The most apical region of the IMC, called the apical cap, is formed by a single vesicle and was shown to participate to the apical complex stability (Back et al. 2020; O’Shaughnessy et al. 2020; Tosetti et al. 2020) Members of the coccidian subgroup possess an additional open-barrel shaped structure made of unique fibers of tubulin polymers termed the conoid (Hu et al. 2002), with two pre-conoidal rings (PCRs) at the top and two short intra-conoidal microtubules on the inside (Dos Santos Pacheco et al. 2020). The conoid lays between the apical polar ring (APR) which serves as microtubule organizing center (MTOC) for the spiraling SPMTs. The apical cap proteins 9 and 10 (AC9, AC10) were previously shown to be SPN resident proteins whose depletion resulted in the loss of the conoid and the APR, and disorganization of the SPMTs at the apical pole (Back *et al.* 2020; Tosetti *et al.* 2020). The conoid is the site of convergence for calcium and lipid-mediated signaling cascades that coordinate microneme secretion and actin polymerization, and is also the site of for the glideosome, the molecular machinery powering parasite gliding motility (Bisio & Soldati-Favre 2019). Strikingly AC9 and AC10 depleted parasites were severely impaired in microneme secretion and consequently unable to move, invade or egress from host cells. We postulated that the loss of the conoid and SPMTs disorganization, could result from the disappearance of the APR. Concordantly another study reported that the double knock-out of two APR proteins (APR1 and KinA) led to the fragmentation of the APR, partial detachment of the conoid and reduced microneme exocytosis (Leung et al. 2017). In contrast, depletion of the mitogen-activated protein kinase (MAPK) ERK7 reportedly cause the loss of conoid in mature parasite without seemingly affecting microneme secretion nor APR integrity (O’Shaughnessy *et al.* 2020). Moreover, AC9 was assigned a dual role in localizing ERK7 at the apical cap and in regulating its kinase activity and substrate specificity (Back *et al.* 2020). Another protein, the conoidal ankyrin repeat-containing protein (CPH1) was also reported to contribute to the conoid integrity in extracellular parasites. CPH1 depleted parasites harbored shortened, partially collapsed conoids, again without seemingly impacting on microneme secretion (Long et al. 2017a).

Here, we have revisited the role of ERK7 and CPH1 in microneme secretion. Depletion of ERK7 phenocopies strictly the depletion AC9 or AC10, leading not only to the loss of the conoid but also the disappearance of the APR and a severe impairment in induced microneme secretion. A comparative proteomics analysis of wild-type and ERK7 or AC9 depleted parasites highlighted the loss of known, as well as novel candidate proteins composing the apical complex. Conversely the accumulation of microneme proteins reflected the severe defect in exocytosis of these secretory organelles.

## Results

### ERK7 is essential for the stability of the conoid complex and the organization of the subpellicular microtubules in mature parasites

Three mitogen-activated protein kinases (MAPKs) are encoded in the genome of *T. gondii* and conserved across the superphylum of the Alveolata, except for TgMAPK-like 1 which is missing in haemosporidia and piroplasmida orders (Lacey et al. 2007; Talevich et al. 2011); their respective localization and function in *T. gondii* and *Plasmodium* spp. are summarized in Figure S1A. Briefly, TgMAPK-like 1 and TgMAPK2 are both involved in different steps of parasite replication (Suvorova et al. 2015; Hu et al. 2020), while TgERK7 is involved in conoid biogenesis thus hampering parasites invasion, egress and dissemination (O’Shaughnessy *et al.* 2020). The *Plasmodium* spp. orthologues of TgERK7 (Pf/PbMAP-1) and TgMAPK2 (PF/PbMAP-2) are dispensable in the asexual blood stage with only Pf/PbMAP-2 required for male gametogenesis (Khan et al. 2005; Rangarajan et al. 2005; Tewari et al. 2005; Hitz et al. 2020).

TgERK7 was carboxy-terminally tagged with the mAID-HA (ERK7-mAID-HA) at the endogenous locus and shown to be tightly downregulated upon addition of auxin (IAA) (Figure S1B) and localized at the apical cap of both mature and daughter cells (Figure S1C). As previously reported, parasites depleted in ERK7 fail to form lysis plaques in a monolayer of human foreskin fibroblasts (HFFs) after 7 days of IAA treatment (Figure S1D). Given the close relationship reported between AC9 and ERK7, the cytoskeletal integrity was checked by deoxycholate (DOC) extraction. As observed in absence of AC9 or AC10, the cytoskeleton of parasites depleted in ERK7 is disassembled (Figure 1A). In addition to the loss of conoid, the APR has disappeared, leading to a wide apical opening, explaining the detached SPMTs visible by DOC extraction (Figure 1B). Ultrastructure expansion microscopy (U-ExM) was applied here for the first time on intracellular parasites to compare with observations made on extracellular parasites, using the ERK7-mAID-HA strain further modified by Ty-epitope tagging of AC2, a marker of the alveolin network (ERK7-mAID-HA / AC2-Ty). As previously shown with AC9 deleted parasites, the absence of ERK7 led to the loss of both conoid and APR in mature parasites, while the nascent daughter cells still possessed an intact apical complex (Figure 1C). Overall AC2 was not dramatically affected by ERK7 depletion; however, occasionally, AC2 was either not present apically or was markedly reduced in some parasites (Figure 1C). The same observation was made with another apical cap protein, ISP1 (Figure S1E).

**Figure 1.**
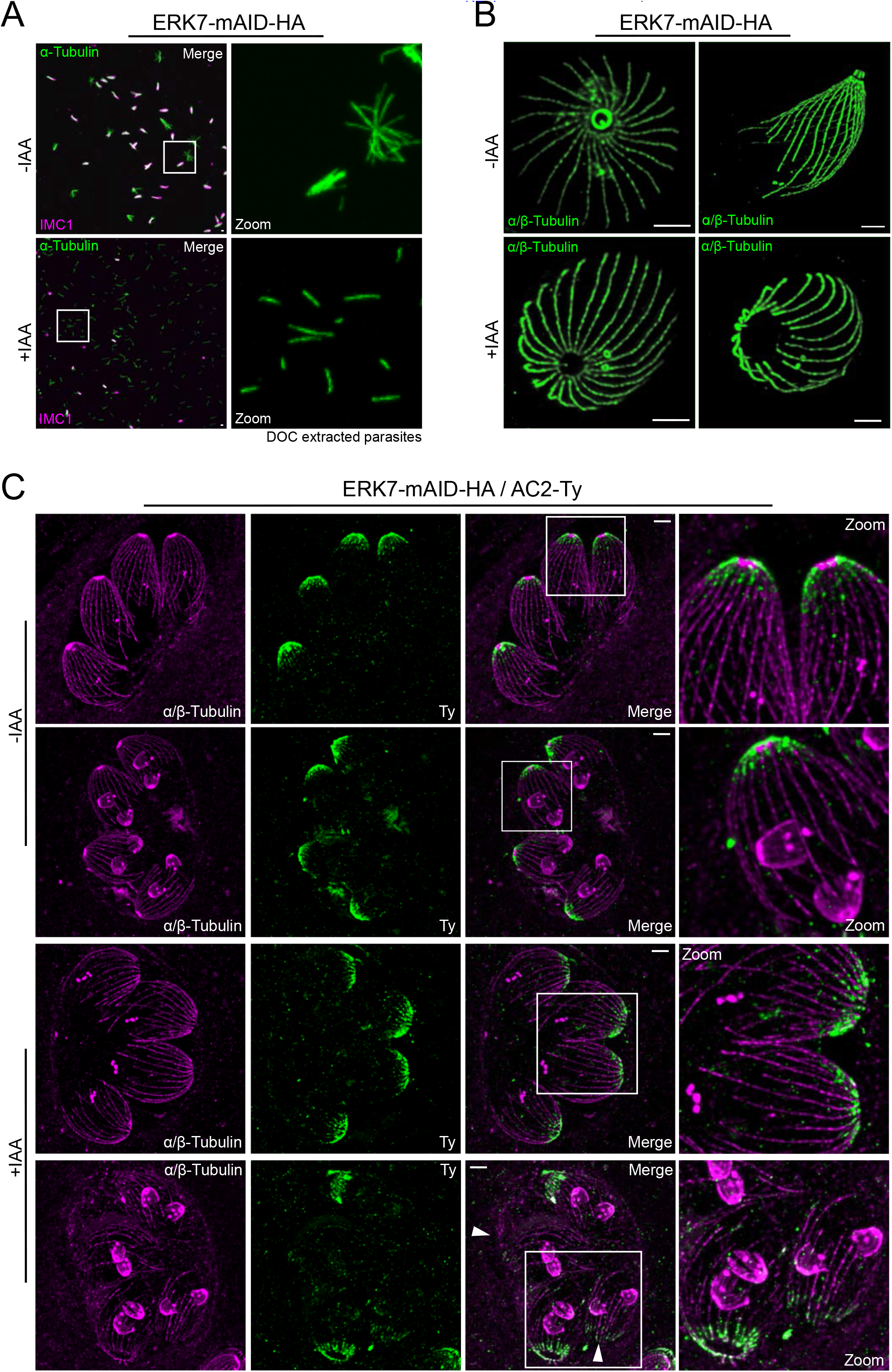
ERK7 depletion caused major cytoskeletal defects at the apical pole. **(A)** Extracellular parasites were extracted with deoxycholate (DOC) and placed on gelatin coverslips. Depletion of ERK7 caused the collapse of the microtubular cytoskeleton as only single microtubules can be visualized by IFA with α-tubulin antibody suggesting that the APR is no more present in those mutants. **(B)** Ultrastructure expansion microscopy (U-ExM) highlighted the absence of the APR and consequently an enlargement of the apical pole in ERK7 depleted parasites. **(C)** U-ExM was applied in intracellular conditions with AC2 tagged in the ERK7 inducible strain. Upon addition of IAA, the APR and conoid are missing exclusively in the mother cell and in some severe cases of SPMTs disorganization the AC2 staining disappears from the apical cap too (arrowhead). Scale bars = 2 mm.

### The apical polar ring is lost in mature ERK7 depleted parasites

To further confirm the loss of the APR, three APR markers were epitope tagged in the ERK7-mAID-HA strain, namely RNG1, APR1 and KinA (Tran et al. 2010; Leung *et al.* 2017). RNG1, a very late maker of parasite division, was not visible at the APR of mature parasites (Figure 2A and 2B; Figure S2A); whereas the protein was still detectable by western blot analysis (Figure 2A). Remarkably, the remaining RNG1 signal is mis-localized in the absence of ERK7, accumulating mainly at the posterior pole of the parasites (Figure 2A; Figure S2A). The two other APR markers were lost exclusively in the mature parasite but present in nascent daughter cells, confirming ERK7 role occurring late in parasite division (Figure 2C and 2D). In contrast to RNG1, APR1 and KinA are both degraded based on western blot analysis (Figure 2C and 2D).

**Figure 2.**
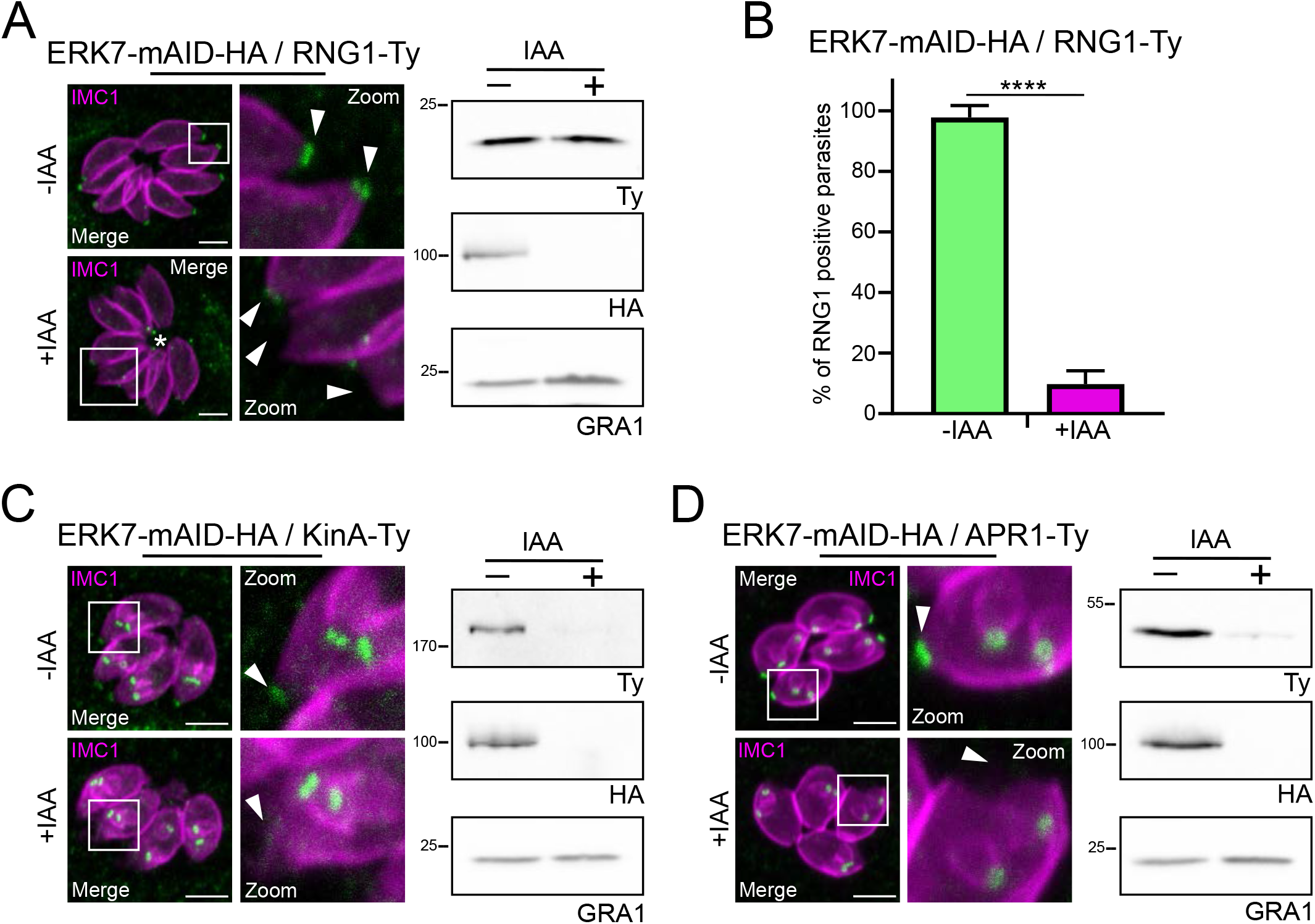
Loss of apical markers in ERK7 depleted parasites. **(A)** RNG1, a late marker of parasite division, failed to be incorporated in most of the parasites. APR and RNG1 signals could be detected in parasite cytoplasm/residual body (asterisk). Western blot analysis confirmed that RNG1 is not degraded. **(B)** Quantification of apical RNG1 in ERK7 depleted parasites. **(C)** and **(D)** APR markers, APR1 and KinA, are incorporated very early during daughter cells formation and are exclusively lost in mature parasites following deletion of ERK7. Western blot analysis of extracellular parasites (no or very few daughter cells present) treated with IAA showed that both markers are degraded. Arrowheads indicate mother cell apex. Scale bars = 2 mm.

### Rhoptries are disorganized in mature ERK7 depleted parasites by NHS-Ester staining coupled to U-ExM

Next, to gain further insight into the morphology of the ERK7 depleted parasites, we coupled U-ExM newly adapted to *T. gondii* (Tosetti *et al.* 2020) with the fluorescent dye N-hydroxysuccinimide ester (NHS-Ester) able to bind to primary amines on proteins. Remarkably, this combination allows the observation of a wide range of anatomical features while approaching the resolution of electron microscopy (EM) (M’Saad & Bewersdorf 2020). This technique allowed the visualization a wide spectrum of recognizable ultrastructures (movie 1 and 2) with a non-exhaustive list of such features presented in individual stacks (Figure S3). As expected, in ERK7 depleted parasites the conoid was clearly absent from mature parasites while still present in forming daughter cells (Figure 3A, movie 3 and 4). Strikingly the neck of the rhoptries also detected with anti-RON9 antibodies, are still apical but dispersed compared to their apical positioning in presence of ERK7. Interestingly, the so-called “neck” of the rhoptries seems to not extend all the way up to the conoid. This observation was also confirmed with an anti-RON4 antibody (data not shown). Furthermore as previously observed by EM for AC9 depleted parasites (Tosetti *et al.* 2020), in the absence of ERK7 the plasma membrane was dilated at the apical pole (Figure 3A).

**Figure 3.**
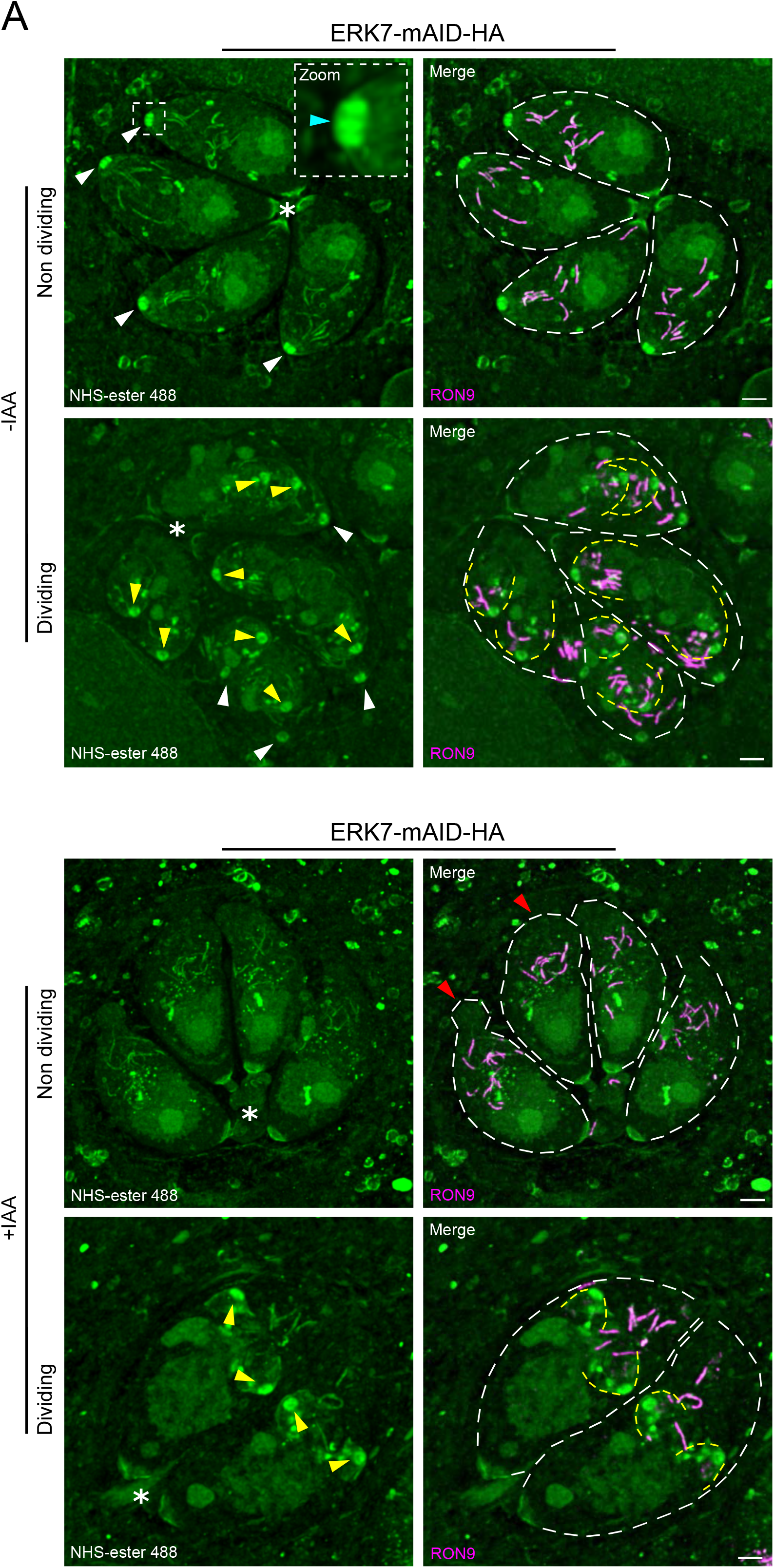
NHS-ester coupled with U-ExM. **(A)** Maximum projection pictures of ERK7 untreated and treated parasites (non-dividing and dividing) from videos 1, 2, 3 and 4. White arrowheads indicate mother cells conoid (blue arrowhead: intraconoidal microtubules); yellow arrowheads highlight daughter cells conoid; red arrowheads indicate deformation of the apical plasma membrane. Scale bars = 2 mm.

### ERK7 depleted parasites are defective in induced microneme secretion

The dilated plasma membrane was examined by staining for the major surface antigen (SAG1) and both in AC9 and ERK7 depleted conditions, the integrity of the membrane appeared compromised in some intracellular parasites (Figure 4A). This leads to an apparent releases of few micronemes outside the parasite body (Figure 4B; Figure S4A and S4B), as previously reported by EM in the absence of AC9 (Tosetti *et al.* 2020).

**Figure 4.**
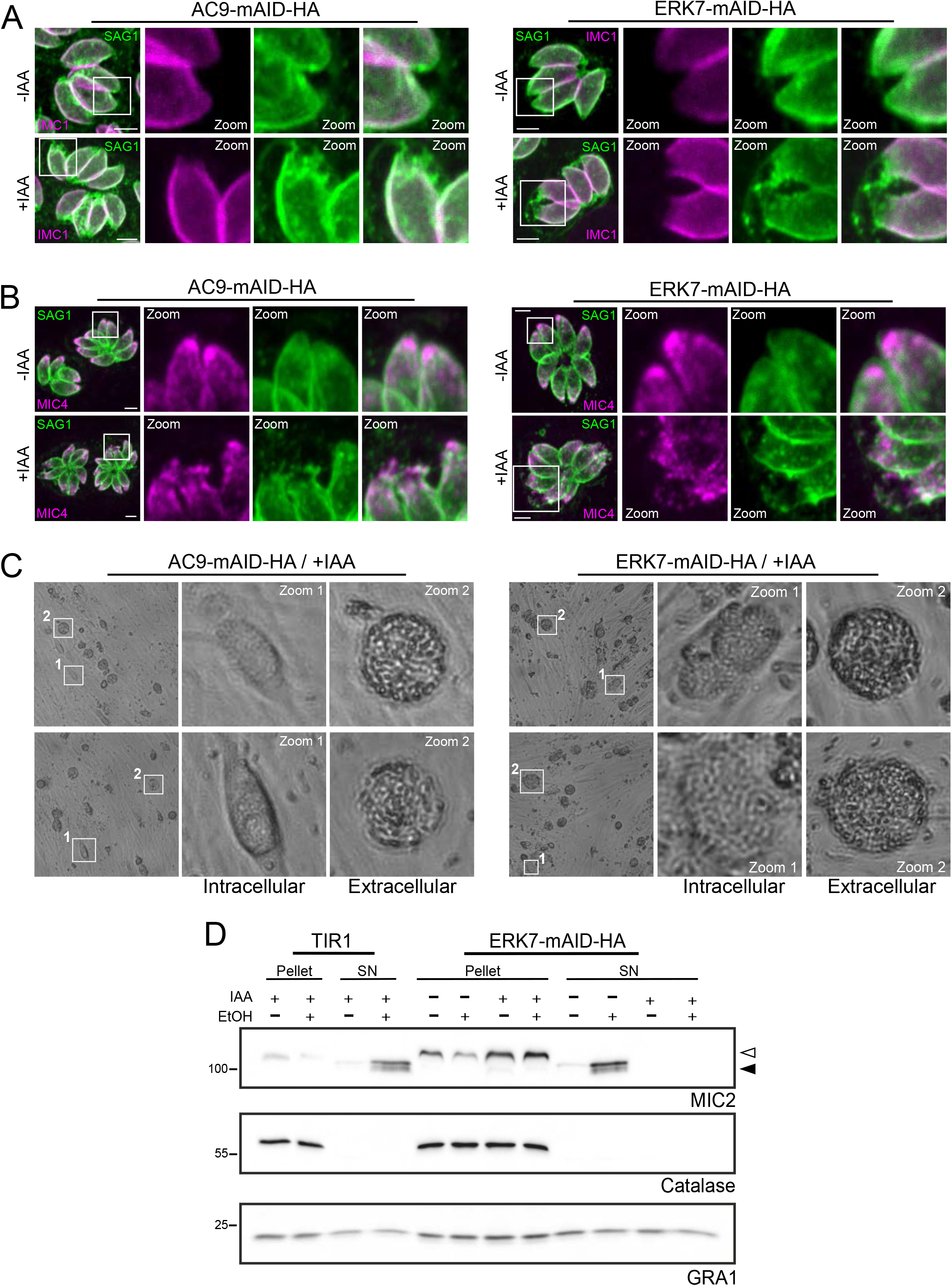
ERK7 depleted parasites are defective in PM integrity and microneme secretion. **(A)** In both AC9 and ERK7 depletion, some parasites per vacuoles showed a defect in plasma membrane integrity at the apical tip highlighted by SAG1 staining and **(B)** leaking of micronemes (MIC4 staining). **(C)** Large bright field images of AC9 and EKR7 parasites treated for 48h with IAA. Depleted parasites showed no defects in intracellular growth; however, parasites remained trapped inside the floating host cell remnant suggesting a microneme secretion impairment. **(D)** Depletion of ERK7 caused a dramatic failure in parasite to secrete microneme when stimulated with ethanol (EtOH). Anti-MIC2 antibodies were used for secretion (white arrow: full length MIC2; black arrow: secreted MIC2), anti-catalase (CAT) to assess parasites lysis and anti-dense granule 1 (GRA1) for constitutive secretion; both pellets and supernatants (SN) were analyzed. Scale bars = 2 μm.

Although AC9 and ERK7 are both essential for parasite survival, depleted mutants are not affected in intracellular growth and replication (Figure 4C). In contrast, both mutants are seemingly able to lyse the parasitophorous vacuole membrane (PVM) but remained trapped inside the host cell even after the host cell detached in floating “bubbles” suggesting at least a partial microneme secretion defect (Figure 4C). While depletion of ERK7 was reported not to impact on microneme secretion, a severe exocytosis defect was observed in absence of AC9 (Back *et al.* 2020; O’Shaughnessy *et al.* 2020; Tosetti *et al.* 2020). The microneme secretion used by O’Shaughnessy *et al,* was based on a luciferase assay applied previously to CPH1-mAID (Long et al. 2017). Here the induced secretion assay was monitored by detection of cleaved released MIC2 in the supernatant and quantified by western blot. Under these conditions, ERK7-mAID-HA treated with IAA showed a complete block in microneme secretion as reported for AC9 (Figure 4D). As control, the CPH1-mAID strain was generated (Figure S4C and S4D) and shown to exhibit a milder defect in microneme secretion indicating that the discrepancy can be explained only in part by the type of assay used (Figure S4E).

### Comparative proteomics of wild-type and ERK7/AC9 depleted parasites identifies novel candidate components of the apical complex

Given the dramatic disappearance of the structure as well as the protein markers of the conoid and APR in both AC9 and ERK7 depleted parasites, we performed a comparative proteomic analysis by mass spectrometry to gain further knowledge about the composition of the apical complex. The datasets analysis revealed a good overall coverage in comparison to the hyperLOPIT dataset (3,832 *T. gondii* detected proteins) (Barylyuk et al. 2020), with a total of 3477 and 3483 proteins detected in ERK7 and AC9 samples, respectively (Figure 5C). In agreement with previously published data and proteomics datasets, numerous known apical complex proteins disappeared in both AC9 and ERK7 depleted parasites (Figure 5A, 5B, 5D and Figure 6A). Out of the 17 most significantly depleted proteins in both datasets, 14 were already localized at the apical pole of the parasite (Figure 6A). Overall, the analysis points toward at least a dozen of plausible novel candidate proteins potentially associated to the apical complex.

**Figure 5.**
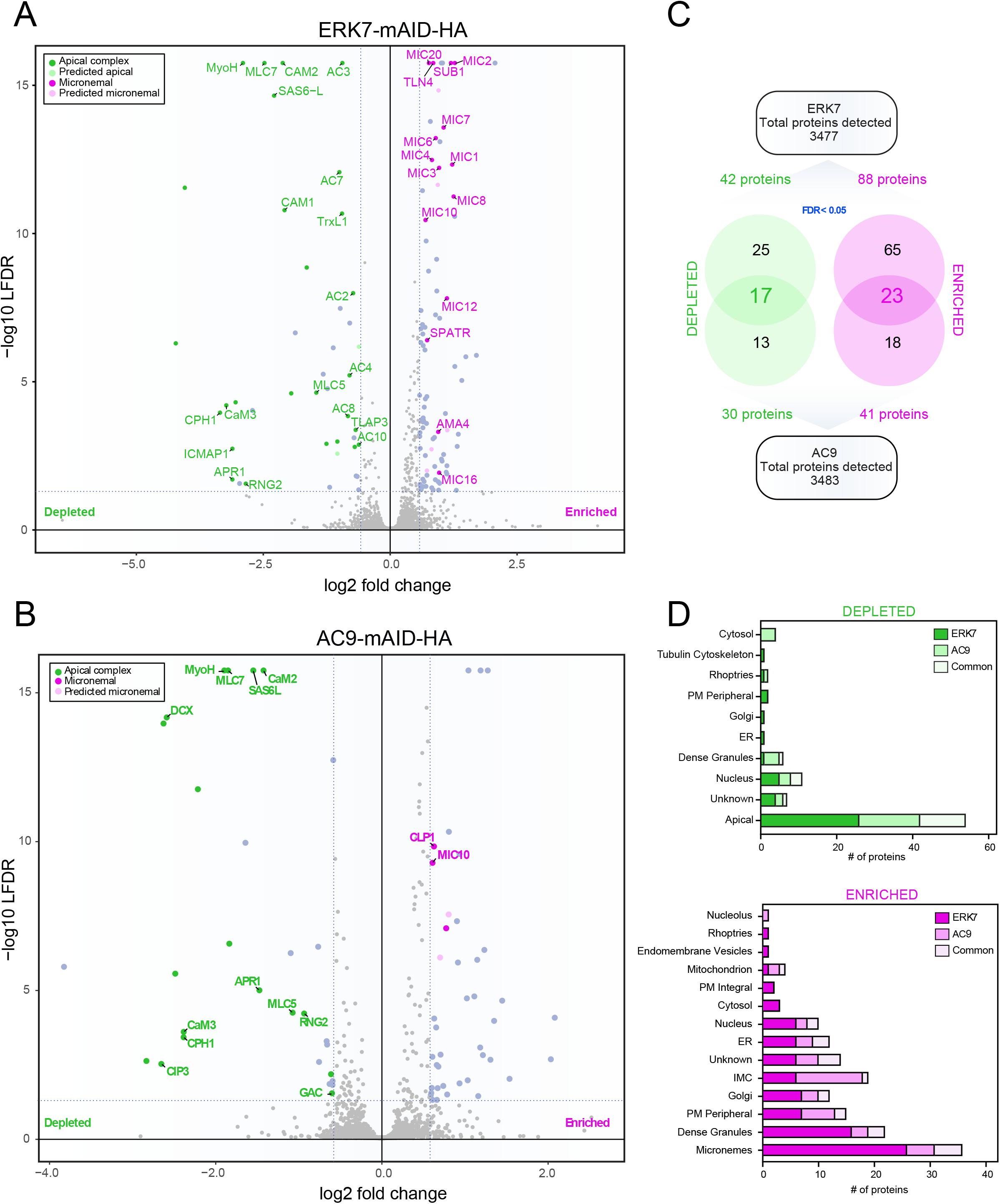
Comparative proteomics of ERK7 and AC9 depleted parasites. **(A)** and **(B)** Volcano plots showing all the proteins found in ERK7 (A) and AC9 (B) mAID parasites and their differential abundance in auxin-treated parasites vs. untreated parasites. The most significant changes are colored and grouped based on their known and predicted (LOPIT) localization. On the left (in green) are proteins that are depleted and that majorly localize to the apical complex. The proteins on the right (in magenta) are the one that are enriched and are found majorly within the micronemes. **(C)** Venn diagrams showing the significant changes in ERK7 and AC9 (proteins individually depleted or enriched) and the common proteins depleted or enriched in both ERK7 and AC9 auxin-treated parasites. **(D)** Predicted localization of enriched and depleted proteins in both ERK7 and AC9 according to hyperLOPIT.

**Figure 6.**
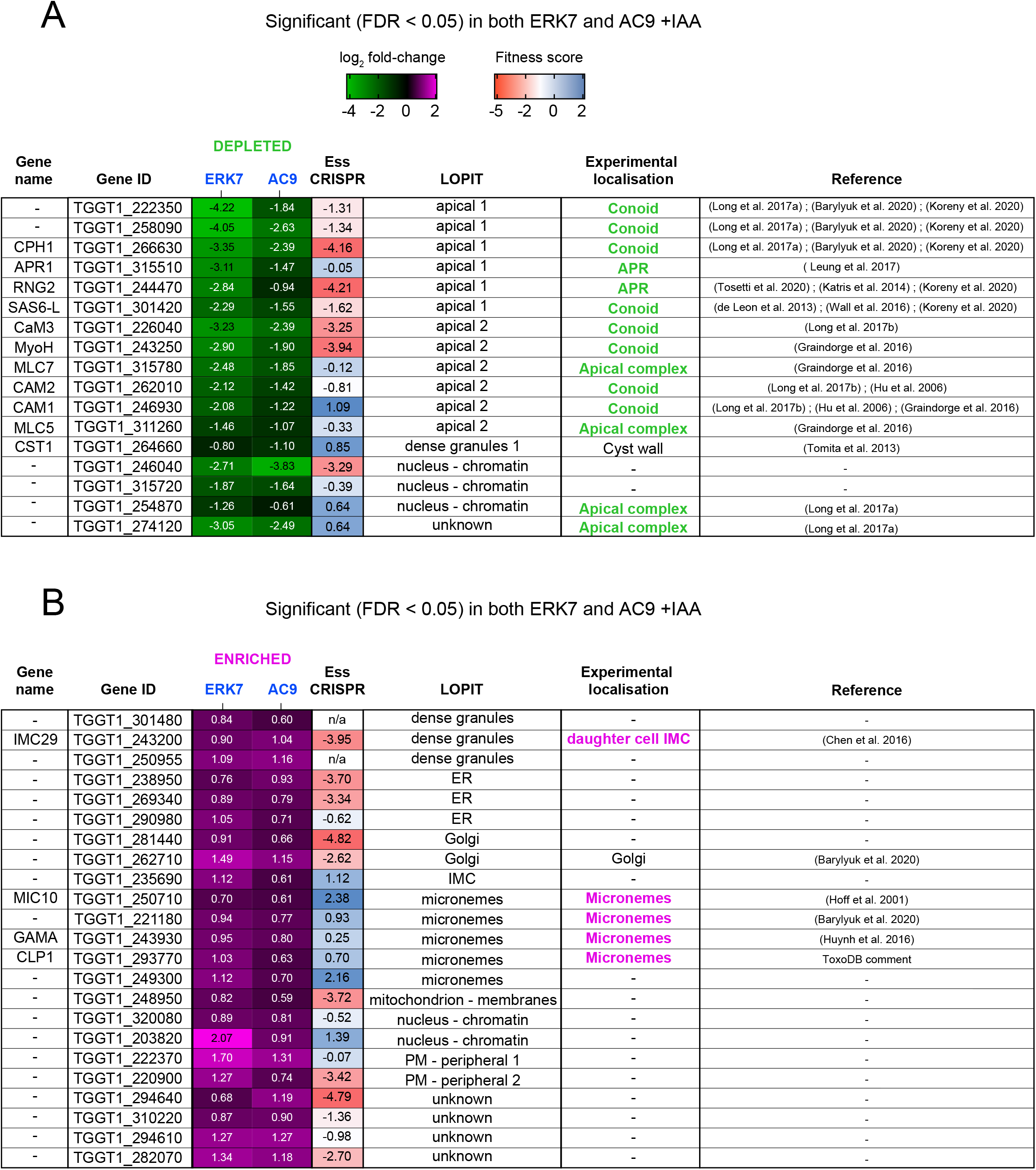
Statistically significant proteins depleted/enriched in both strains. **(A)** and **(B)** Heat map showing the most significant changes in both ERK7 and AC9 (auxin-treated) conditions. In green (A) are the proteins that are depleted and in magenta (B) are the proteins that are enriched. Predicted (LOPIT) localizations and previously localized proteins are also indicated.. (Hoff et al. 2001; Hu et al. 2006; de Leon et al. 2013; Tomita et al. 2013; Katris et al. 2014; Graindorge et al. 2016; Huynh & Carruthers 2016; Wall et al. 2016; Chen et al. 2017; Leung *et al.* 2017; Long *et al.* 2017a; Long et al. 2017b; Barylyuk *et al.* 2020; Tosetti *et al.* 2020)

### Comparative proteomics of wild-type and ERK7 and AC9 depleted parasites show an accumulation of microneme proteins and components of the IMC

Beside the loss of components of apical complex and conversely, other distinct sets of proteins accumulate in ERK7 and AC9 depleted parasites, respectively (Figure 5A, 5B, 5D and Figure 6B). In the absence of ERK7, known microneme proteins as well as proteins predicted by hyperLOPIT (Barylyuk et al. 2020) to be associated with micronemes are significantly more abundant. This is easily accounted for by the severe microneme exocytosis defect observed in the mutant parasites. In AC9 depleted parasites, microneme proteins also accumulated however other proteins predicted to be localized at the IMC or dense granules are overrepresented. Out of the 17 proteins enriched in both datasets and not localized experimentally to date, 7 share a cell cycle transcription profile common to known IMC proteins (Figure 6B and Figure S5). A comprehensive list of the comparative proteomics analysis of ERK7 and AC9 mutants is presented in Dataset 1.

## Discussion

AC9 is a component of the alveolin network proposed to serve as regulatory platform for ERK7 kinase activity (Back *et al.* 2020). In this study we have revisited the role of ERK7 in the integrity of the apical complex of *T. gondii* in light of unmatching phenotypic consequences compared to AC9 depletion (O’Shaughnessy *et al.* 2020; Tosetti *et al.* 2020). ERK7 depleted parasites phenocopied AC9 and AC10 depletion except that APR was shown to be still present and microneme secretion was not affected. Here a new ERK7-mAID-HA strain was generated and confirmed to be severely affected in invasion and egress from infected cells. U-ExM analysis proved that like AC9 and AC10, ERK7 is required to maintain the conoid stability but also to preserve the tethering of the SPMTs in mature parasites. In contrast to the previous report, the depletion of ERK7 leads the disappearance of the APR with the degradation of APR1 and KinA and the mislocalization of RNG1. As a late marker of the APR, RNG1 might not be a structural component *per se* of the APR but rather associated to the structure only in mature parasites. The absence of the APR might explain why this protein end up in the residual body in ERK7 depleted parasites. The absence of APR also explains the wide apical opening observed in U-ExM and why detached individual microtubules are observed in deoxycholate extraction experiments. The APR is likely lost upon parasite final maturation as it presumably serves as MTOC for the SPMTs.

The NHS-Ester staining coupled with U-ExM offers novel opportunities to study in detail the internal structures of the parasite in its vacuolar niche. For example, in the less well-studied parasites like *Cryptosporidium spp.* in which a very sparse number of antibodies have been raised for localization studies (Wilke et al. 2018). The NHS-Ester could be a staining of great value as it can virtually reveal all structures within the cell. Also, as shown in this study, some structures are very well stained with this method. The “ring structures” systematically observed presumably corresponds to the micropore or the ring structures described with GAPM2 and GAPM3, two resident proteins of the IMC (Nichols et al. 1994; Harding et al. 2019). The basal complex of the parasite appears extremely dense with this technique, probably highlighting a density of protein complexes present at the basal pole of the parasite. In addition, very small structures in dividing parasites like the centrocone and centromeres are surprisingly evident with the NHS-Ester staining while only EM could achieve such level of details (Francia et al. 2020). Further improvements of these methods could potentially help retaining membrane or lipid-bound epitope as well as reaching greater expansion ratio without the need of heavy instruments (Damstra et al. 2021).

The induced microneme secretion monitored here by excretory-secretory antigen detection via western blot provides radically opposing outcome for ERK7-mAID-HA compared to the published results based on the luciferase assay. The presence of the conoid seems absolutely required for microneme secretion as no processed MIC2 can be detected in supernatant from parasites depleted in ERK7, AC9 or AC10 (Tosetti *et al.* 2020). Since O’Shaughnessy *et al,* used a very sensitive assay based on MIC2 fused to luciferase reporter we also generated a CPH1-mAID mutant previously analyzed by the same method to include in the analysis (Long *et al.* 2017a). The two assays are rather concordant for CPH1 depleted parasites although a mild defect in microneme secretion was not detected in the luciferase-based assay (Long et al. 2017). The accumulation of further processing of MIC2, only visible by western blot-based assay, reflects a defect in gliding motility due to the lack of posterior translocation of MIC2, and hence the over-representation of MIC2 protein trimmed by the apical subtilisin-like serin protease SUB1 (Lagal et al. 2010). It is plausible that the luciferase assay is too sensitive for the detection of minor defects microneme exocytosis, yet ERK7 depleted parasites have a very strong defect. Given that a fraction of these mutants was shown to exhibit a dilated apical plasma membrane, they might be more prone to lysis and could perturb the assay if parasites are not carefully washed before the experiment. In a conceptually distinct assay, O’Shaughnessy et al, also assessed microneme secretion by monitoring mVenus diffusion from the PV to the host cell cytoplasm. This diffusion is indicative of a permeabilization of the PVM by the action of the secreted microneme protein perforin PLP1 (Kafsack et al. 2009). AC9 and ERK7 depleted parasites are indeed able to lyse the PVM but remained trapped inside the host cell suggesting that a low level of microneme exocytosis was occurring in a few intracellular parasites but not sufficient to trigger complete egress and hence resulted in parasites trapped in floating detached host cells. Alternatively the discrepancies observed in ERK7 mutant with regard to the disappearance of the APR and ability to secrete microneme could be explained by revertant parasites that fail to respond anymore to IAA as mentioned by the authors (O’Shaughnessy *et al.* 2020).

Capitalizing on the loss of APR and conoid that resulted in degradation of several apical complex protein in both ERK7/AC9 depleted strains, we embarked on a proteomics analysis to compare the mutants to wild-type parasites and identify novel components of the affected structures. The comparative analysis between each replicate revealed a striking quantitative reproducibility for the vast majority of proteins. Consequently, the subsets of proteins disappearing or accumulating in the samples emerged very clearly. A large set of proteins already known or predicted to be associated with the apical complex were detected while only a small number of potential new apical complex proteins were found, potentially reflecting some technical limitation of mass spectrometry in detecting small, low abundant or not easily digested proteins. Alternatively, the combination of approaches used so far to determine the repertoire of apical complex protein might be almost comprehensive.

Of relevance, known, as well as predicted microneme proteins are accumulating in ERK7 and AC9 deleted parasites, concordantly with their impairment in microneme exocytosis. This tendency was more prominent in the ERK7 depleted parasites since twice as many proteins were found significantly enriched in those parasites compared to the AC9 depleted ones (88 versus 41). This might be due to the intrinsic nature and role of the two proteins: while AC9 is a structural component of the alveolin network, ERK7 is an active kinase (Back *et al.* 2020). Some proteins enriched in the absence of ERK7 could be affected by the absence of kinase activity and not only be a consequence of the loss APR and conoid and the disassembly of SPMTs. In the absence of ERK7 or AC9, some proteins predicted by hyperLOPIT to localize at the dense granules, ER, Golgi, nucleus also accumulate (Barylyuk *et al.* 2020). The experimental assessment of their localization will be instrumental as some protein such as the product of TGGT1_254870, mainly predicted to localize at the nucleus was previously localized at the apical pole of the parasite (Long *et al.* 2017). Similarly, the product of TGGT1_243200, predicted to localize to the dense granules is associated to the IMC of nascent daughter cells (Chen *et al.* 2016).

Overall, with novel technologies such as U-ExM, NHS-Ester staining and comparative proteomics mass-spectrometry, a deeper look into the role of ERK7 was taken in this study. Our observations revealed that the loss of ERK7, a kinase likely phosphorylating several components (known and unknown) of the apical complex, indeed dislocates the conoid complex. The severe microneme secretion defect also strengthens previous observations that exocytosis of these organelles, and as a consequence, proper egress, gliding, motility and invasion of intracellular parasites, might not be achieved without the presence of a structurally intact conoid.

## Abbreviations

AC: apical cap protein
SPMTs: subpellicular microtubules
IMC: inner membrane complex
ERK7: Extracellular signal-regulated kinase 7

## Supplementary materials

**Figure S1.**
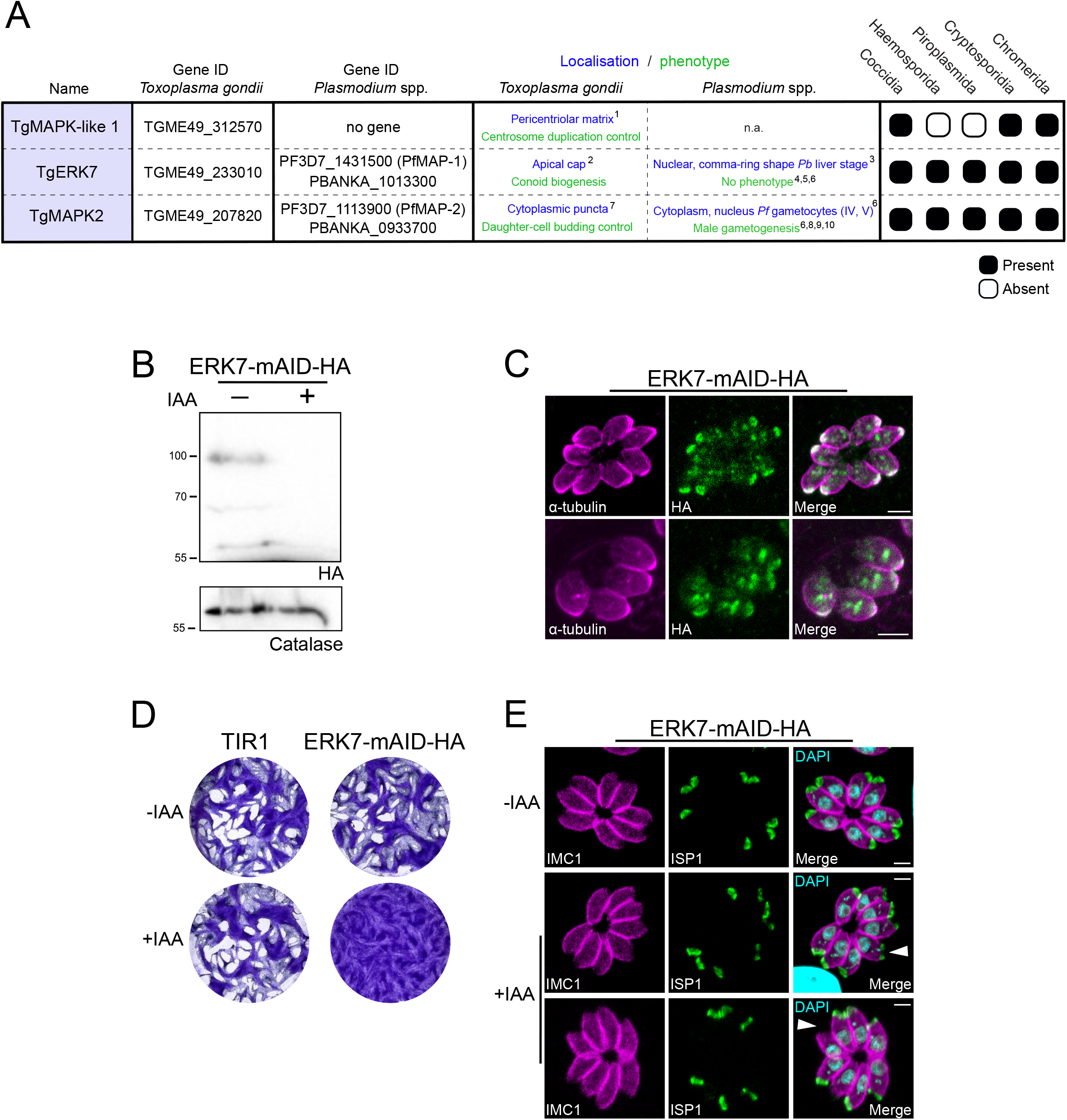
**(A**) Mitogen-activated protein kinases (MAPKs) present in *T. gondii* and *Plasmodium* spp. **1** (Suvorova *et al.* 2015), **2** (O’Shaughnessy *et al.* 2020), **3** (Wierk et al. 2013), **4** (Dorin-Semblat et al. 2007), **5** (Fang et al. 2018), **6** (Hitz *et al.* 2020), **7** (Hu *et al.* 2020), **8** (Khan *et al.* 2005), **9** (Rangarajan *et al.* 2005), **10** (Tewari *et al.* 2005). **(B)** ERK7 was C-terminally tagged with the mAID-HA construct and was tightly downregulated upon addition of auxin (IAA). **(C)** ERK7 localized at the apical cap of both mature and daughter cells. Like AC9 and AC10, ERK7 is among the earliest markers appearing during endodyogeny, before the daughter cytoskeletal basket. **(D)** ERK7 depleted parasites failed to form lysis plaques after 7 days. TIR1 represents the parental strain. **(E)** Localization of another apical cap protein (ISP1) was not majorly impacted by ERK7 degradation; nevertheless, the apical region of some parasites per vacuoles is clearly enlarged and some shows no or reduced ISP1 signal (arrowhead). Scale bars = 2 μm.

**Figure S2.**
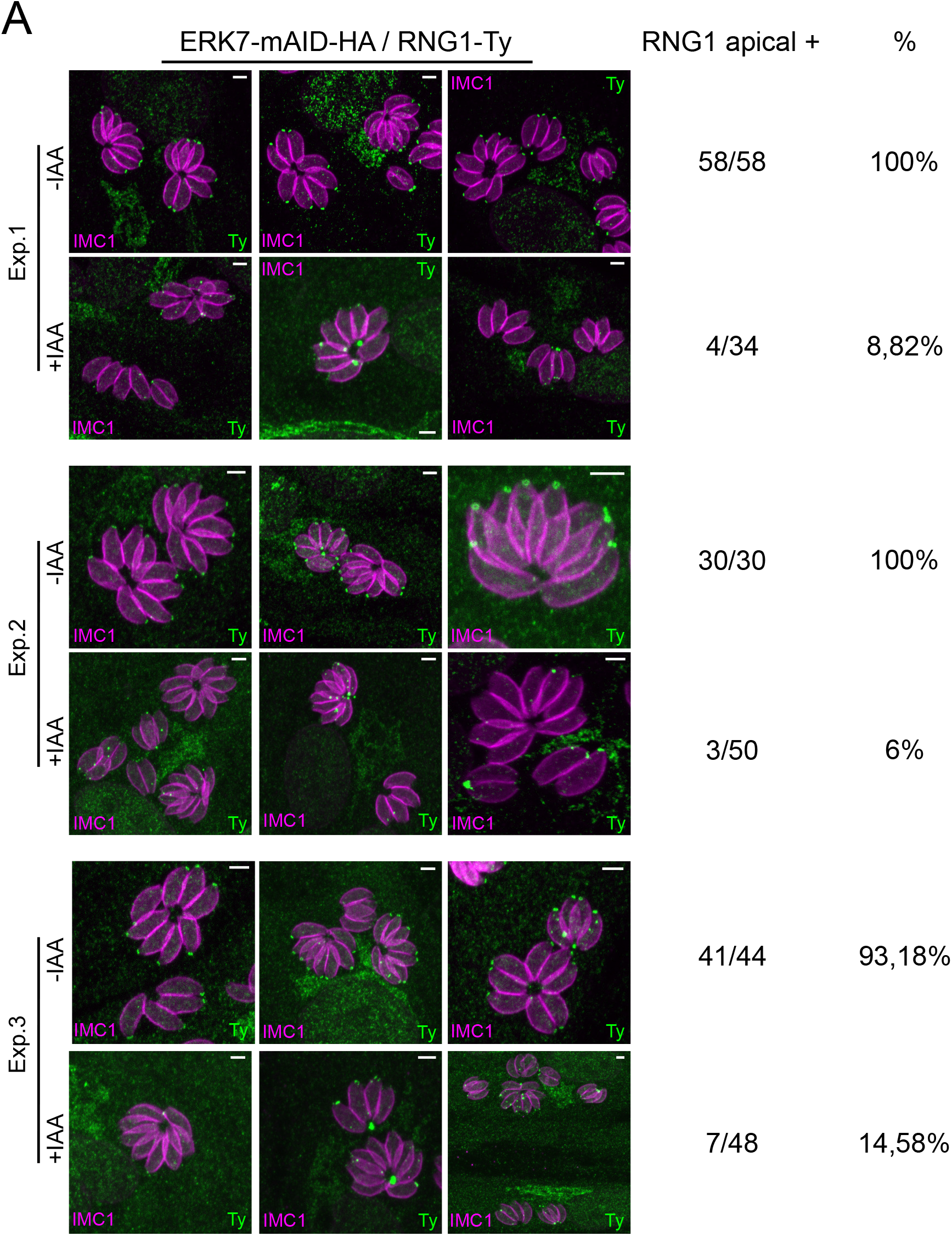
**(A)** Raw pictures and quantification of RNG1 in ERK7 inducible strain. Scale bars = 2 μm.

**Figure S3.**
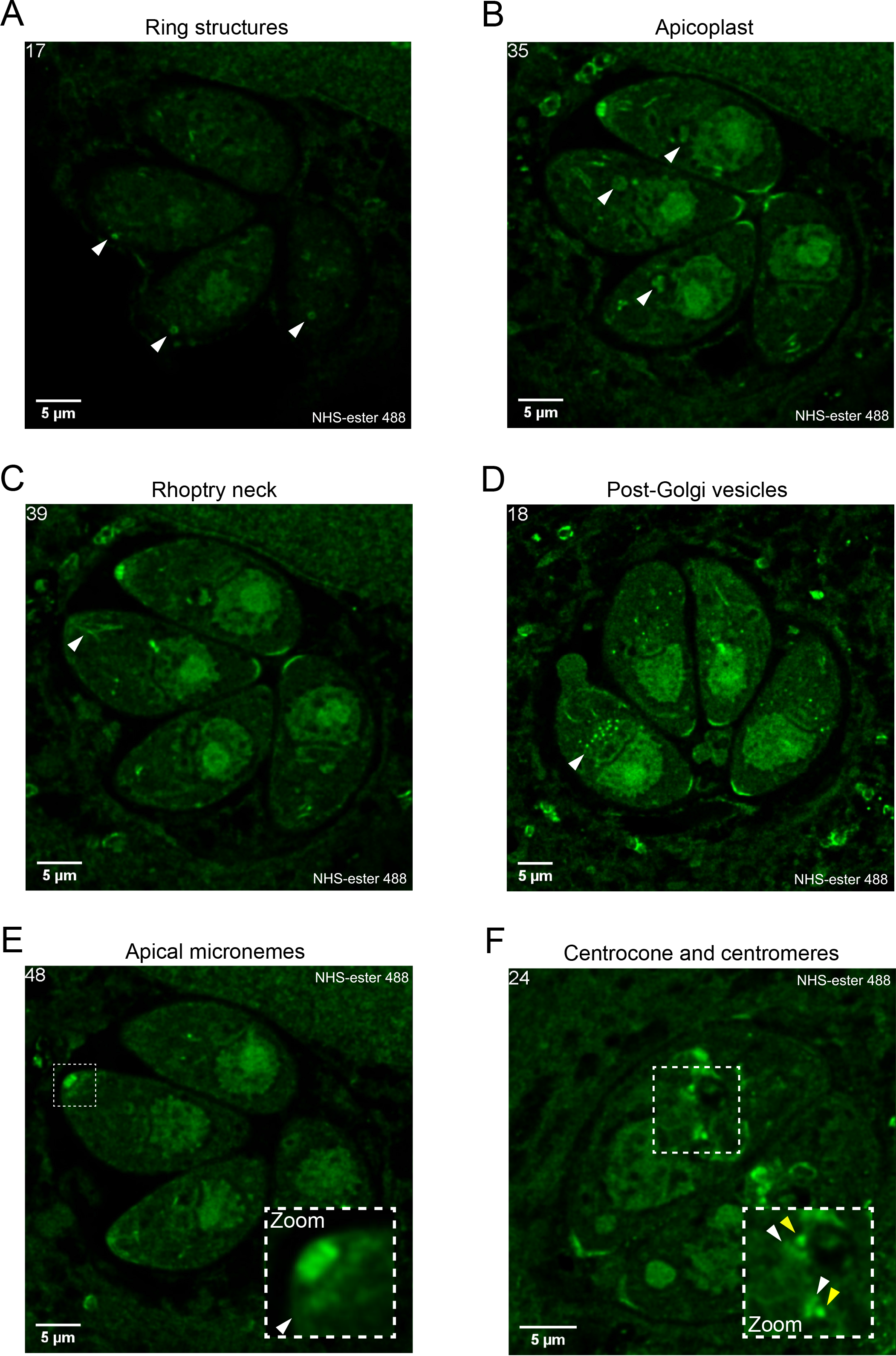
Individual stacks from videos 1, 3 and 4 highlighting the broad spectrum of NHS-ester staining for ultrastructural elements and organelles. On top left the stack number. **(A)** Ring structures observed on parasite surface (from video 1). **(B)** Apicoplast (from video 1). **(C)** Rhoptry neck (from video 1). **(D)** Post-Golgi vesicles (from video 3). **(E)** Apical micronemes collar just below the conoid (from video 1). **(F)** Centrocone (white arrowhead) and centromere (yellow arrowhead) during parasite division (from video 4).

**Figure S4.**
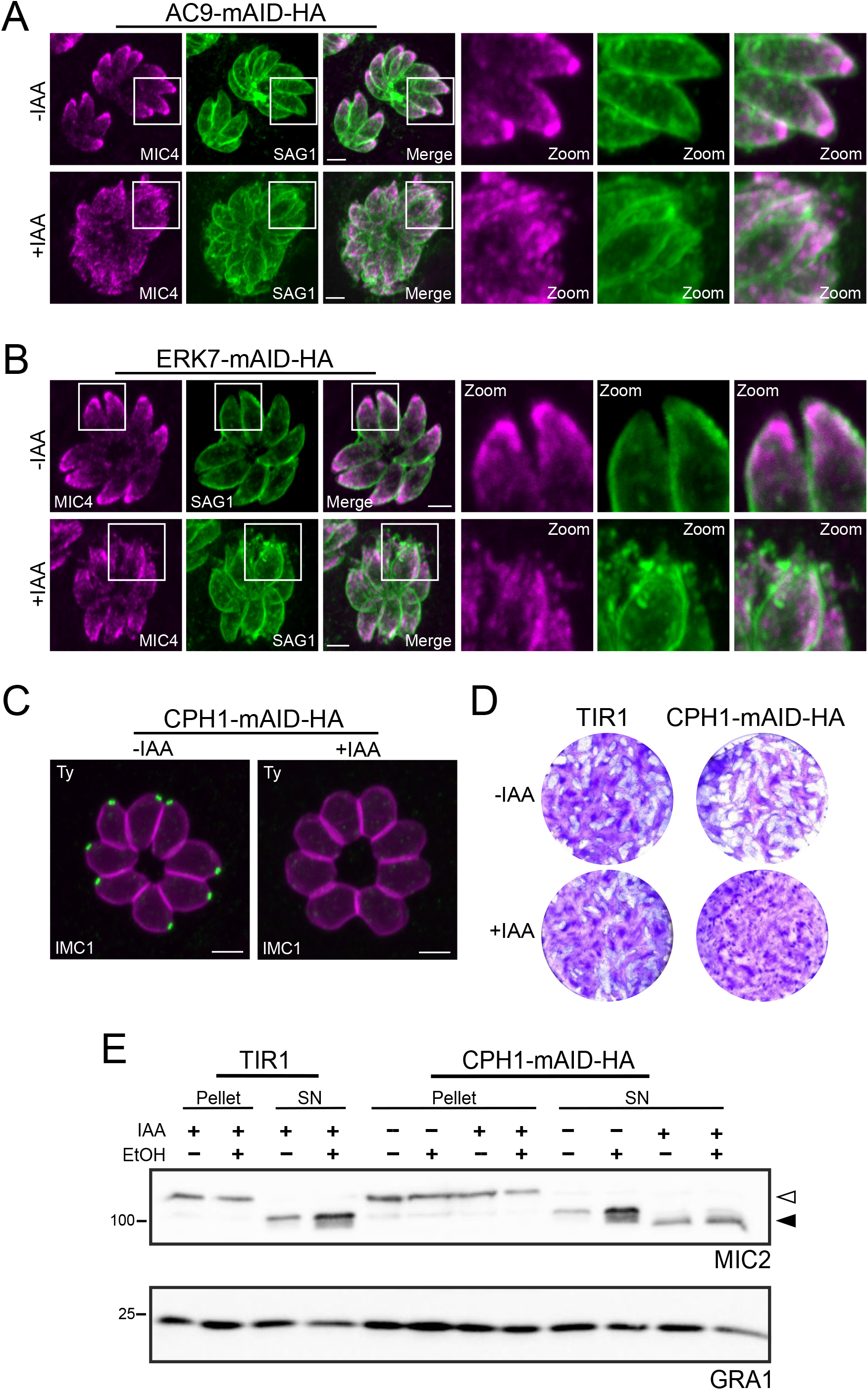
**(A)** and **(B)** Additional pictures of membrane defects in both AC9 and ERK7 and microneme leaking. **(C)** CPH1 was terminally tagged with the mAID-HA construct and was tightly downregulated upon addition of IAA by IFA. **(D)** CPH1 depleted parasites failed to form lysis plaques after 7 days. TIR1 represents the parental strain. **(E)** Depletion of CPH1 caused a moderate defect in microneme secretion when stimulated with ethanol (EtOH). Anti-MIC2 antibodies were used for secretion (white arrow: full length MIC2; black arrow: secreted MIC2) and anti-dense granule 1 (GRA1) for constitutive secretion; both pellets and supernatants (SN) were analyzed. Scale bars = 2 μm.

**Figure S5.**
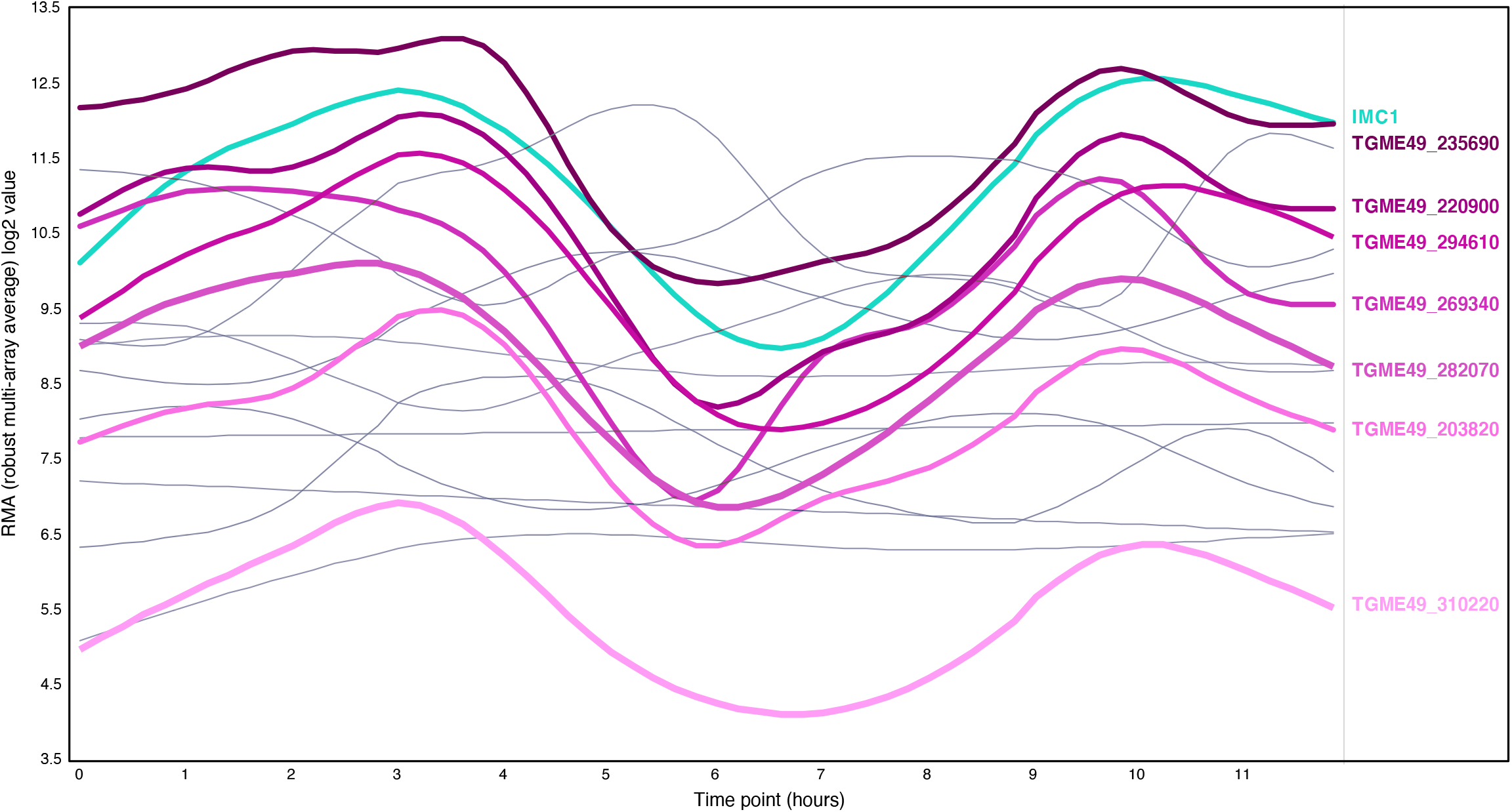
Expression profile of proteins with no “experimental localization” presented in the figure 6B. Out of 17 proteins, 7 of them (thick lines / shades of magenta) share a similar expression profile with a typical IMC protein, namely IMC1 (cyan). The 10 others, with different expression profiles, are presented with thin gray lines. Data is retrieved from the ToxoDB website (Transcriptomic data “T. gondii ME49 Cell Cycle Expression Profiles (RH)” from Behnke *et al*).

**Video 1**

ERK7 untreated parasite (-IAA), non-dividing cells.

**Video 2**

ERK7 untreated parasite (-IAA), dividing cells.

**Video 3**

ERK7 treated parasite (+IAA), non-dividing cells.

**Video 4**

ERK7 treated parasite (+IAA), dividing cells.

## Material and methods

### Generation of strains

*T. gondii* were grown in human foreskin fibroblasts (HFFs, American Type Culture Collection-CRL 1634) in Dulbecco’s Modified Eagle’s Medium (DMEM, Gibco) supplemented with 5% fetal calf serum (FCS), 2 mM glutamine and 25 mg/ml gentamicin. Human Foreskin Fibroblasts (HFF) were obtained from the ATCC (CRL-2088 Ref# CCD1072Sk). ERK7-mAID-HA (TGGT1_233010) was obtained by transfecting a gRNA obtained by Q5 site-directed mutagenesis kit (New England Biolabs) on the pSAG1::Cas9-U6::sgUPRT vector (Shen et al. 2014) with primer 5’-CACGCCGCATTTGTTGACTGGTTTTAGAGCTAGAAATAGC-3’ and a KOD PCR obtained with F primer 5’-TTTCAGTCTGCGTCCAAGACATACAACAGCGCTAGCAAGGGCTCGGG-3’ and R primer 5’-CGGCTCTTCTTTGACACAAAGCAAGAGTCTATACGACTCACTATAGGG-3’ as described in (Brown et al. 2018). Depletion of ERK7-mAID-HA was obtained with 500 mM of auxin (IAA) (Brown et al. 2017). Knock-in of APR1, KinA and RNG1 were performed as previously described (Tosetti *et al.* 2020). Cloning were performed in *E. coli* XL-1 Gold chemo-competent bacteria. Freshly egressed *T. gondii* tachyzoites were transfected by electroporation and mycophenolic acid (25 mg/mL) and xanthine (50 mg/mL) (for HXGPRT cassette) or pyrimethamine (1 mg/ml) (for DHFR cassette) were employed to select resistant parasites.

### Immunofluorescence assay

Parasites were grown in HFF cells with cover slips in 24-well plates for 24–30 hr before fixation with either 4% PFA/0.05% glutaraldehyde (PFA/GA) or cold methanol and neutralized in 0.1M glycine/PBS for 5 min. For SAG1 staining, parasites were permeabilized with 0.1% saponin instead of Triton X-100 which was used for all other IFA. The following antibodies were used: rabbit anti-IMC1 (Frenal et al. 2014), rabbit anti-HA (Sigma), mouse α-acetylated α-tubulin (6-11B-1; Santa Cruz Biotechnology), mouse anti-ISP1 (Beck et al. 2010), rabbit anti-ARO (Mueller et al. 2013), mouse anti-Ty, anti-MIC2 and anti-ROP2-4 (gifts from J-F Dubremetz, Montpellier), anti-tubulin AA344 scFv-S11B (β-tubulin) and AA345 scFv-F2C (α-tubulin), secondary antibodies Alexa Fluor 405-, Alexa Fluor 488-, Alexa Fluor 594-conjugated goat a-mouse/a-rabbit. For western blot, secondary peroxidase conjugated goat a-rabbit or mouse antibodies (Sigma) were used. Confocal images were obtained with a Zeiss laser scanning confocal microscope (LSM700; objective apochromat 63x/1.4 oil). Ultrastructure expansion microscopy (U-ExM) was performed as previously described (Tosetti *et al.* 2020) and images taken with a Leica TCS SP8 STED with lightning deconvolution. For intracellular conditions, the parasites were seeded on non-confluent host cells in order to guarantee the good expansion of the sample. All U-ExM gels were carefully measured to assure a minimum expansion ratio of 4X. The NHS-Ester staining (Thermofisher, DyLight™ 488 NHS Ester) was used at 5μg/mL and incubated 1 hour in PBS. Confocal and expansion images were processed with ImageJ and LAS X, respectively. Images were obtained at the Bioimaging core facility of the Faculty of Medicine at University of Geneva.

### Plaque assay

HFF were inoculated with fresh parasites and grown for 7 days with or without IAA. HFF were then fixed with PAF/GA, washed once with PBS and stained with crystal violet

### Microneme secretion

After ~48 hr of grown parasites were harvested and centrifuged at 1000g. Pellets were then washed twice in intracellular buffer pre-warmed at 37°C (5 mM NaCl, 142 mM KCl, 1 mM MgCl2, 2 mM EGTA, 5.6 mM glucose and 25 mM HEPES, pH 7.2). Next, parasites were incubated at 37°C for 15 min in DMEM containing 2% of ethanol (EtOH). After induction, parasite were centrifuged at 1000 g for 5 min at 4°C and supernatants (SN) collected in different tubes and centrifuged at higher speed (2000 g) for 5 min at 4°C to clean from parasite debris; pellets were washed once in PBS. Pellets and SNs were analyzed using anti-MIC2, anti-catalase (CAT; parasite lysis control) and anti-dense granule 1 (GRA1; constitutive secretion control) antibodies by western blot.

### Deoxycholate extraction

Freshly egressed parasites (−/+ IAA) were deposited on top of poly-L-Lysine-coated coverslips and treated with 10 mM deoxycholate (20 min at room temperature). Parasites were fixed with cold methanol for 8 min and immunofluorescence performed with α-acetylated α-tubulin antibody.

### Mass spectrometry

#### Sample preparation

Parasite pellets were resuspended in 100 μl of 1 % RapiGest Surfactant (Waters) in 0.1M triethylammonium bicarbonate (TEAB) and 7.5 mM Dithioerythritol (DTE). Samples were heated for 5 min at 95°C. Lysis was performed by sonication (6 x 30 sec.) at 70% amplitude and 0.5 pulse. Samples were kept 30 sec. on ice between each cycle of sonication. Samples were centrifuged for 5 min. at 16’000 x g. Protein concentration was measured by Bradford assay and 50 μg of each sample was subjected to protein digestion as follow: sample volume was adjusted to 100 μl with 0.1M TEAB to obtain a final concentration of rapigest 0.1%. 2 μl of Dithioerythritol (DTE) 50 mM in distilled water were added and the reduction was carried out at 60°C for 1h. Alkylation was performed by adding 2 μl of iodoacetamide (400 mM in distilled water) during 1 hour at room temperature in the dark. Overnight digestion was performed at 37 °C with 5 μl of freshly prepared trypsin (Promega; 0.2 μg/μl in TEAB 0.1M). To remove RapiGest, samples were acidified with TFA, heated at 37°C for 45 min. and centrifuged 10 min. at 14’000 g. Supernatants were then desalted with a C18 microspin column (Harvard Apparatus, Holliston, MA, USA) according to manufacturer’s instructions, completely dried under speed-vacuum and stored at −20°C.

#### ESI-LC-MSMS

Samples were diluted in 50 μl of loading buffer (5% CH3CN, 0.1% FA) and 2 μl were injected on column. LC-ESI-MS/MS was performed on an Orbitrap Fusion Lumos Tribrid mass spectrometer (Thermo Fisher Scientific) equipped with an Easy nLC1200 liquid chromatography system (Thermo Fisher Scientific). Peptides were trapped on a Acclaim pepmap100, C18, 3μm, 75μm x 20mm nano trap-column (Thermo Fisher Scientific) and separated on a 75 μm x 500 mm, C18 ReproSil-Pur (Dr. Maisch GmBH), 1.9 μm, 100 Å, home-made column. The analytical separation was run for 180 min using a gradient of H2O/FA 99.9%/0.1% (solvent A) and CH3CN/FA 99.9%/0.1% (solvent B). The gradient was run from 5 % B to 28 % B in 160 min, then to 40% B in 20 min, then to 95%B in 10 min with a final stay of 20 min at 95 % B. Flow rate was of 250 nL/min an total run time was of 210 min. Data-dependent analysis (DDA) was performed with MS1 full scan at a resolution of 120’000 FWHM followed by as many subsequent MS2 scans on selected precursors as possible within 3 second maximum cycle time

#### Database search

Peak lists (MGF file format) were generated from raw data using the MS Convert conversion tool from ProteoWizard. The peaklist files were searched against the *Toxoplasma gondii GT1* database (ToxoDB, release 48, 8460 entries) combined with an in-house database of common contaminant using Mascot (Matrix Science, London, UK; version 2.5.1). Trypsin was selected as the enzyme, with one potential missed cleavage. Precursor ion tolerance was set to 10 ppm and fragment ion tolerance to 0.02 Da. Variable amino acid modifications were oxidized methionine and deaminated (NQ). Fixed amino acid modification was carbamidomethyl cysteine. The Mascot searches were validated using Scaffold 4.11.1 (Proteome Software). Peptide identifications were accepted if they could be established at greater than 91.0% probability to achieve an FDR less than 0.1% by the Scaffold Local FDR algorithm. Protein identifications were accepted if they could be established at greater than 96.0% probability to achieve an FDR less than 1.0% and contained at least 2 identified peptides. Protein probabilities were assigned by the Protein Prophet algorithm (Nesvizhskii et al. 2003). Proteins that contained similar peptides and could not be differentiated based on MS/MS analysis alone were grouped to satisfy the principles of parsimony.

#### Comparative proteomics

Bioinformatics analysis: For the comparative proteomics analysis, the average of the three replicates (minus and plus auxin, IAA) was used to compute the false discovery rate (FDR). A cutoff of 0.05 on the FDR was applied to obtain the most significant hits. For the volcano plots, the log10 of the FDR (y-axis) and the log2 fold change (FC), comparing minus and plus auxin treated parasites (x-axis) was computed. A cutoff of 0.5 was applied to the log2FC to mark the significant changes. The plots were generated using R statistical software, and the heat map with the common significant changes (depleted and enriched proteins) in both ERK7 and AC9 plus auxin was generated using GraphPad Prism v9. All the raw data can be found in Supplementary Table S1.

## Acknowledgements

We thank the Bioimaging Core Facility and the Proteomic Core Facility (University of Geneva – CMU) for their excellent technical support. We thank the Brochet lab and the Guichard/Hamel lab for their helpful discussion to improve the U-ExM technology. This work was supported by the Swiss National Science Foundation (SNSF) 310030_185325 to DSF.

